# Phylogenetic and population structure analyses uncover pervasive misclassification and help assessing the biosafety of *Pseudomonas alloputida* for biotechnological applications

**DOI:** 10.1101/2020.12.22.423983

**Authors:** Hemanoel Passarelli-Araujo, Sarah H. Jacobs, Glória R. Franco, Thiago M. Venancio

**Author notes:** Corresponding authors Av. Alberto Lamego 2000, P5 sala 217; Parque Califórnia, Campos dos Goytacazes, RJ, Brazil, CEP: 28013-602, HPA; TMV.

## Abstract

The *Pseudomonas putida* group comprises strains with biotechnological and clinical relevance. *P. alloputida* was proposed as a new species and highlighted the misclassification of *P. putida*. Nevertheless, the population structure of *P. alloputida* remained unexplored. We retrieved 11,025 *Pseudomonas* genomes and used *P. alloputida* Kh7^T^ to delineate the species. The *P. alloputida* population structure comprises at least 7 clonal complexes (CCs). Clinical isolates are mainly found in CC4 and acquired resistance genes are present at low frequency in plasmids. Virulence profiles support the potential of CC7 members to outcompete other plant or human pathogens through a type VI secretion system. Finally, we found that horizontal gene transfer had an important role in shaping the ability of *P. alloputida* to bioremediate aromatic compounds such as toluene. Our results provide the grounds to understand *P. alloputida* genetic diversity and safety for environmental applications.

## 1. Introduction

The genus *Pseudomonas* is diverse and composed of Gram-negative bacteria [1, 2]. Since the first description of *Pseudomonas*, in 1894, the taxonomy of the genus has evolved in parallel with new methodologies to discriminate species [1]. The *Pseudomonas* taxonomy was inconsistent until the division in rRNA homology groups [2]. Nevertheless, because of the limited resolution of the 16S rRNA gene for intraspecific classification, other housekeeping genes were later used to improve taxonomic resolution [3, 4].

The *Pseudomonas putida* group includes other species, such as *P. monteilii*, *P. fulva*, *P. plecoglossicida*, and *P. putida sensu stricto*. Species from this group are ubiquitous in soil and water, and several strains have been isolated from various environments, such as polluted soils and plant roots [5, 6]. *P. putida* species are well known to perform many functions such as plant growth promotion, bioremediation, and protection against plant pathogens [7].

Because of its high metabolic diversity, applications of *P. putida* have been proposed for environmental, industrial, and agricultural uses [5]. Among well-known *P. putida* species, *P. putida* KT2440 is a model organism to study plant-microbe interactions [8, 9]. KT2440 was isolated from garden soil in Japan and has become a workhorse for industrial biotechnology because of its high metabolic versatility, genetic accessibility, and stress-resistance [10]. In addition to the biotechnological application of *P. putida* species, some clinical isolates have been described as reservoirs of resistance genes [11, 12], raising concerns about its biosafety.

Recently, Keshavarz-Tohid et al. (2019) showed that *P. putida* KT2440 and other known *P. putida* strains (e.g. BIRD-1, F1, and DOT-T1E) are distant from the type strain *P. putida* NBRC 14164^T^ and hence should be classified as a member of a novel species, *Pseudomonas alloputida*, whose type strain is Kh7 (=CFBP 8484^T^ =LMG 29756^T^) [13]. Here, we report the population structure of *P. alloputida*, which was used to estimate the diversity and distribution of bioremediation and plant growth promotion genes. Further, we used the inferred population structure to better understand the prevalence of antibiotic resistance and virulence genes and to assess, in detail, the potential biosafety concerns regarding this species.

## 2. Results and Discussion

### 2.1. Phylogeny and classification of *Pseudomonas alloputida*

We obtained 11,025 *Pseudomonas* genomes available in RefSeq in June 2020, out of which 10,457 had completeness greater than 90% according to BUSCO [14]. We computed the pairwise distances between each isolate using mashtree [15] to compute the distance tree of the genus, which is highly diverse (Figure 1). We mapped each genome deposited in the NCBI RefSeq as *P. putida* in the tree and found that the highest density of genomes falls within a monophyletic group of 439 isolates with average nucleotide identity (ANI) values between 84% and 100% (Figure S1a); this clade corresponds to the *P. putida* group (Figure 1) and comprises other species such as *P. plecoglossicida*, *P. monteilli*, and *P. fulva*.

**Figure 1.**
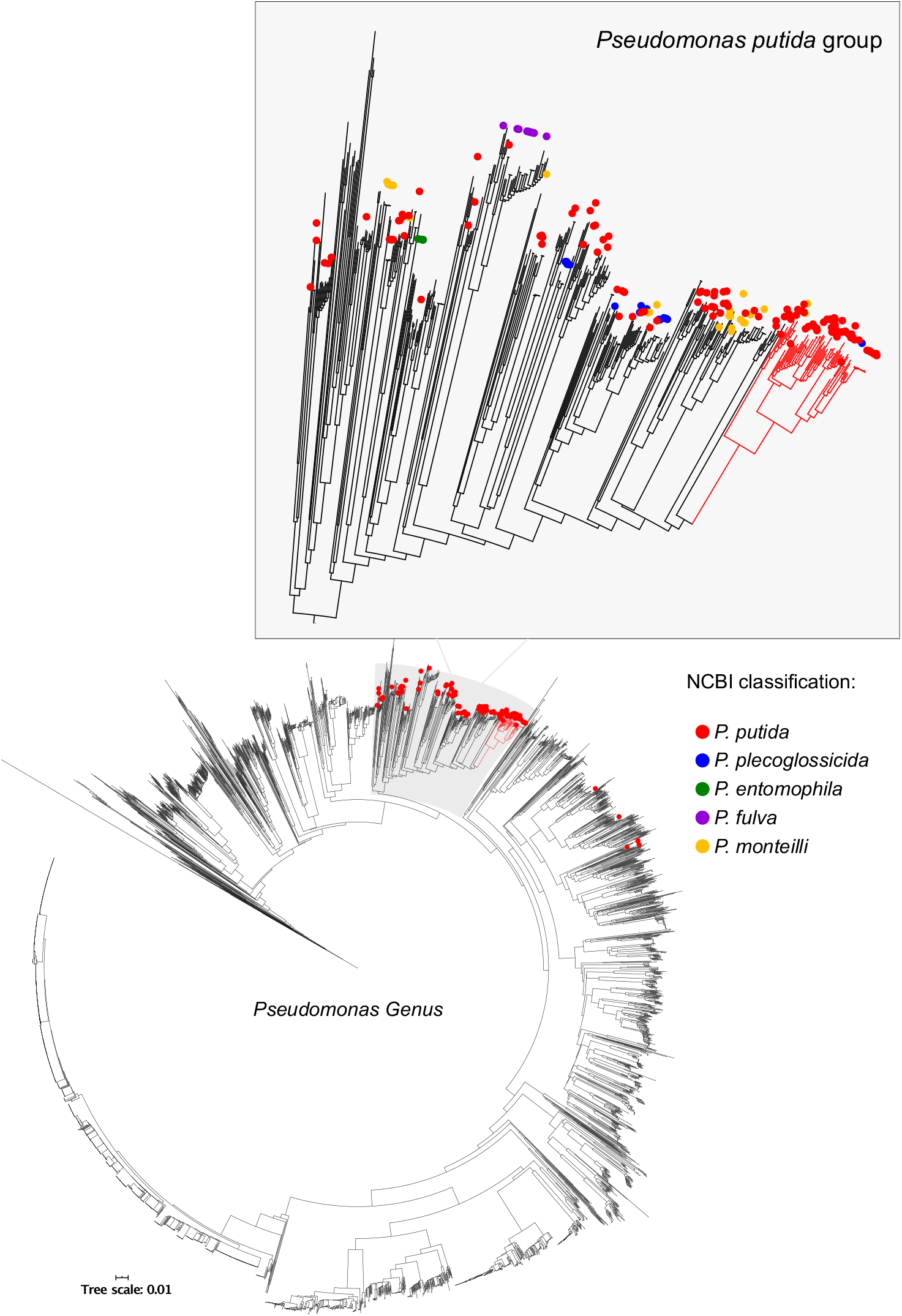
Distance tree of 10,457 *Pseudomonas* genomes from *Pseudomonas* genus. Genomes classified as *P. putida* according to NCBI are marked as red circle. *P. putida* group was highlighted to assess the distribution of misclassified genomes. *P. alloputida* branches are colored in red.

ANI analysis provides a raw estimate of bacterial species [16]. A minimum threshold of 95% ANI has been used to attain species membership, a value that has been empirically defined based on correlations with DNA-DNA hybridization and 16S rRNA thresholds [16, 17]. We used *P. alloputida* Kh7^T^ as an anchor-strain to evaluate the ANI values from other isolates in the *P. putida* group. The sorted distribution of ANI values from Kh7^T^ showed an abrupt break around 95%, supporting its effectiveness as a threshold to delineate *P. alloputida* (Figure S1b). The isolate previously classified as *P. monteilli* IOFA19 (GCA_000633915.1) had the lowest ANI value within the predicted species (95.48%), followed by a drop to 91.13% (Figure S1b).

We conducted a network analysis to assess the species composition according to ANI > 95% with members from the *P. putida* group. We observed a discrete number of cohesive clusters that would correspond to the expected number of species within the *P. putida* group (Figure 2). We retrieved other species such as *P. asiatica*, *P. soli*, and *P. monteilli*, as well as some potentially novel species (Table S1). In this analysis, a species was defined as a cluster containing a type strain and at least three genomes, or a cluster without a type strain, but with at least ten connected genomes. We retrieved the main species from the *P. putida* group and found clusters that most likely correspond to new species (Figure 2). Further, the species number is likely underestimated, as new genomes would increase the number of connections in the network.

**Figure 2.**
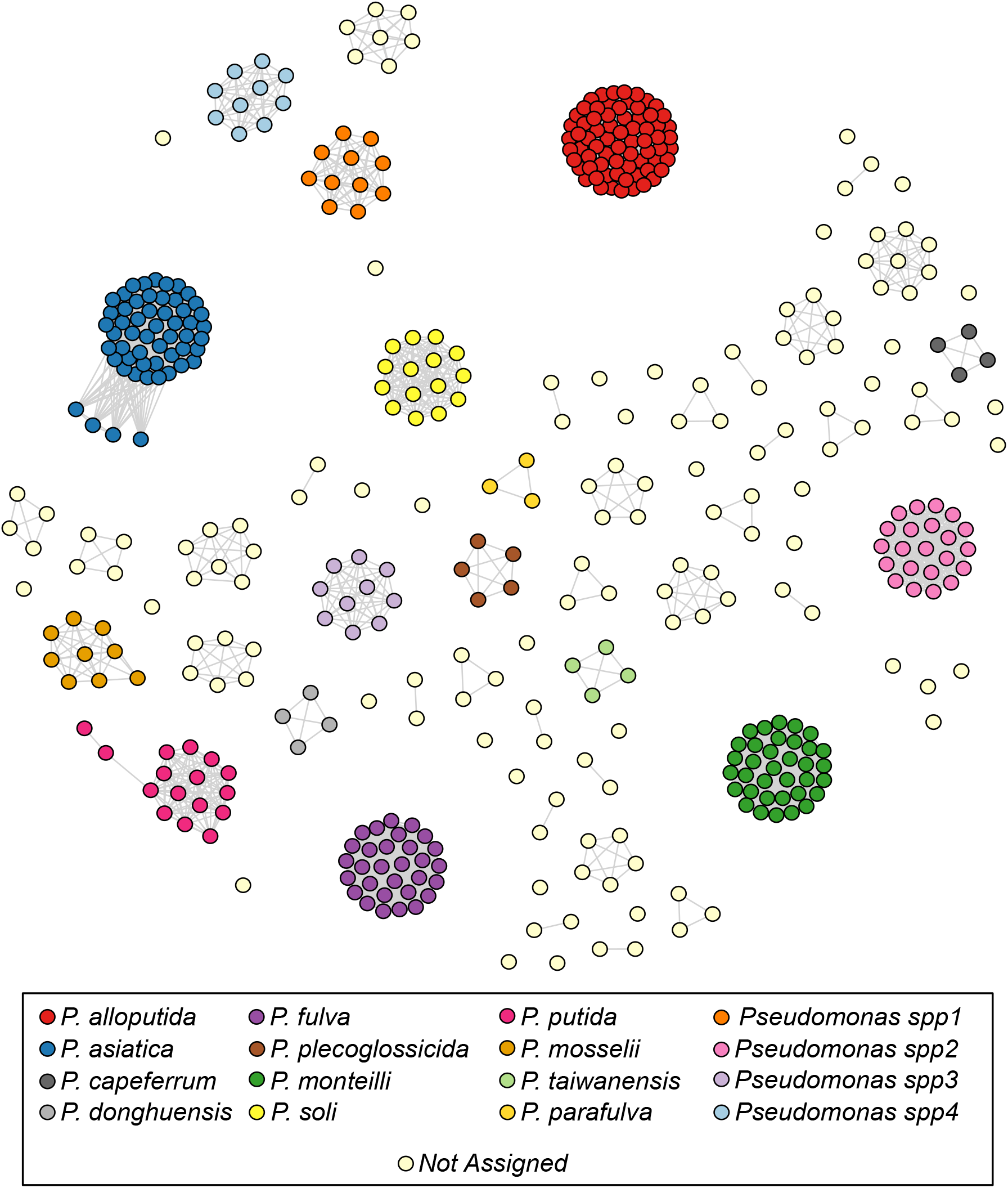
Network analysis of isolates from the *P. putida* group. Nodes represent isolates and edges connect isolates with at least 95% of average nucleotide identity. Clusters with type strains or at least ten genomes were highlighted, as they represent either known or potentially novel species.

The poor classification of *P. putida* isolates is a subject of concern. We observed a clear separation of the clusters with type strains for *P. putida* (NBRC 14164^T^) and *P. alloputida* (Kh7^T^) (Figure 2). *P. putida* and *P. alloputida* comprise groups with 16 and 68 genomes, respectively (Figure 2, Table S1). The greater number of *P. alloputida* genomes might have an historical explanation. Although the phylogenetic separation of NBRC 14164^T^ from other main *P. putida* strains has been noticed before [18], KT2440, a *P. alloputida* isolate [13], has also been used to categorize *P. putida* genomes over the years [6, 8, 18].

The use of NBRC 14164^T^ to delimit *P. putida sensu stricto* highlights that many well-known *P. putida* genomes belong to other species. Isolates that are well known for their ability to promote plant growth (W618) [19], to oxidize manganese (GB-1) [20], and to damage human tissues (HB3267) [11], are neither *P. putida* nor *P. alloputida* strains, as they belong to different groups in the network. For example, HB3267 was classified as *P. putida* because of its close phylogenetic relationship with the nicotine degrader S16 [21]. HB3267, as well as DLL-E4, SF1, and S11, were proposed as members of a new species, *Pseudomonas shirazica* [13]. However, we found that these strains, along with S16, grouped with *Pseudomonas asiatica* type strain, confirming *P. shirazica* as an heterotypic synonym of *P. asiatica* [22]. Henceforth, we focused our analyses in the novel *P. alloputida* species because of its greater number of genomes and of the presence of key strains associated with bioremediation, plant growth promotion, and biocontrol.

### 2.2. Pangenome analysis

An effective way to investigate the evolution of a given population is through pangenome analysis. A pangenome is defined as the total set of genes in a given species [23], which is subdivided into core genes, when present in all isolates; accessory genes, when present in at least two (but not in all) isolates or; exclusive genes. By using 68 isolates, the *P. alloputida* pangenome comprises 25,782 gene families, of which 3803 (14.75%) constitute the core genome (i.e. genes present at least 95% of the isolates). By analyzing the slope of the curve (α= 0.417), we inferred that *P. alloputida* has an open pangenome [23] (Figure S2). Our estimated α value is much lower than the maximum threshold used to define an open pangenome (α < 1), which is in line with a previous study [24]. Further, this low a value implies high rates of new gene families will be found if more isolates are included in the analysis.

The high number of gene families in the *P. alloputida* pangenome is explained by unique and low-frequency genes. Only 6,373 genes families (24.71%) are found between 5% to 95% of the isolates, while 15,606 (60.53%) are found in less than 5% of the isolates, including 10,917 unique genes. The high number of low-frequency genes could be partially attributed to fragmented genomes. However, reference genomes such as DOT-T1E showed 267 unique genes, far more than the average of 160.5 unique genes found across the dataset. Although the number of low-frequency genes may be overestimated, the high prevalence of unique genes among closely-related genomes indicates a high turnover of unstable genes that might be adaptive under transient selective pressures in their environments.

We also estimated the genomic fluidity (φ) of *P. alloputida*. The φ estimator is a robust metric that represents the ratio of unique gene families to the sum of gene families, averaged over randomly chosen genomes pairs [25]. The smaller the φ, the greater the genes shared by a pair of randomly selected genomes. Analyzing φ instead of the core genome proportion provides a more realistic measure of cohesiveness within a species, particularly because low-frequency genes directly affect the pangenome size. *P. alloputida* has φ = 0.20 ± 0.04, indicating that random pairs of *P. alloputida* genomes have an average 20% and 80% of unique and shared genes, respectively.

### 2.3. Population structure

Determining relationships between isolates can provide novel insights into the metabolic diversity of a given species. The Multilocus Sequence Typing (MLST) analysis is a technique to characterize genomes based on single-nucleotide polymorphisms (SNPs) within a few housekeeping genes. MLST schemes are available for several species [26]. In an MLST analysis, each combination of SNPs defines a Sequence Type (ST) that can be linked to form Clonal Complexes (CC) [26]. A variation of classical MLST is the core genome MLST (cgMLST), which provides greater resolution by using SNPs from the entire core genome [27]. Here, we used 225,009 SNPs obtained from the *P. alloputida* core genome to reconstruct the phylogenetic tree and the cgMLST profile.

The cgMLST tree unveiled 7 CCs (Figure 3a), out of which CC1 is the most distant from the rest of the population, a trend that is also supported by the ANI analysis (Figure S3). Six out of the 9 clinical isolates were found in CC4 and three in CC7. We also checked whether the same clustering pattern could be obtained by analyzing the presence/absence patterns of accessory genes. We estimated the Jaccard distance for genes present between 5% − 95% of the isolates to perform a Principal Coordinate Analysis (PCoA), which allowed us to resolve all the main groups, particularly CC1, CC4, and CC7 (Figure 3b).

**Figure 3.**
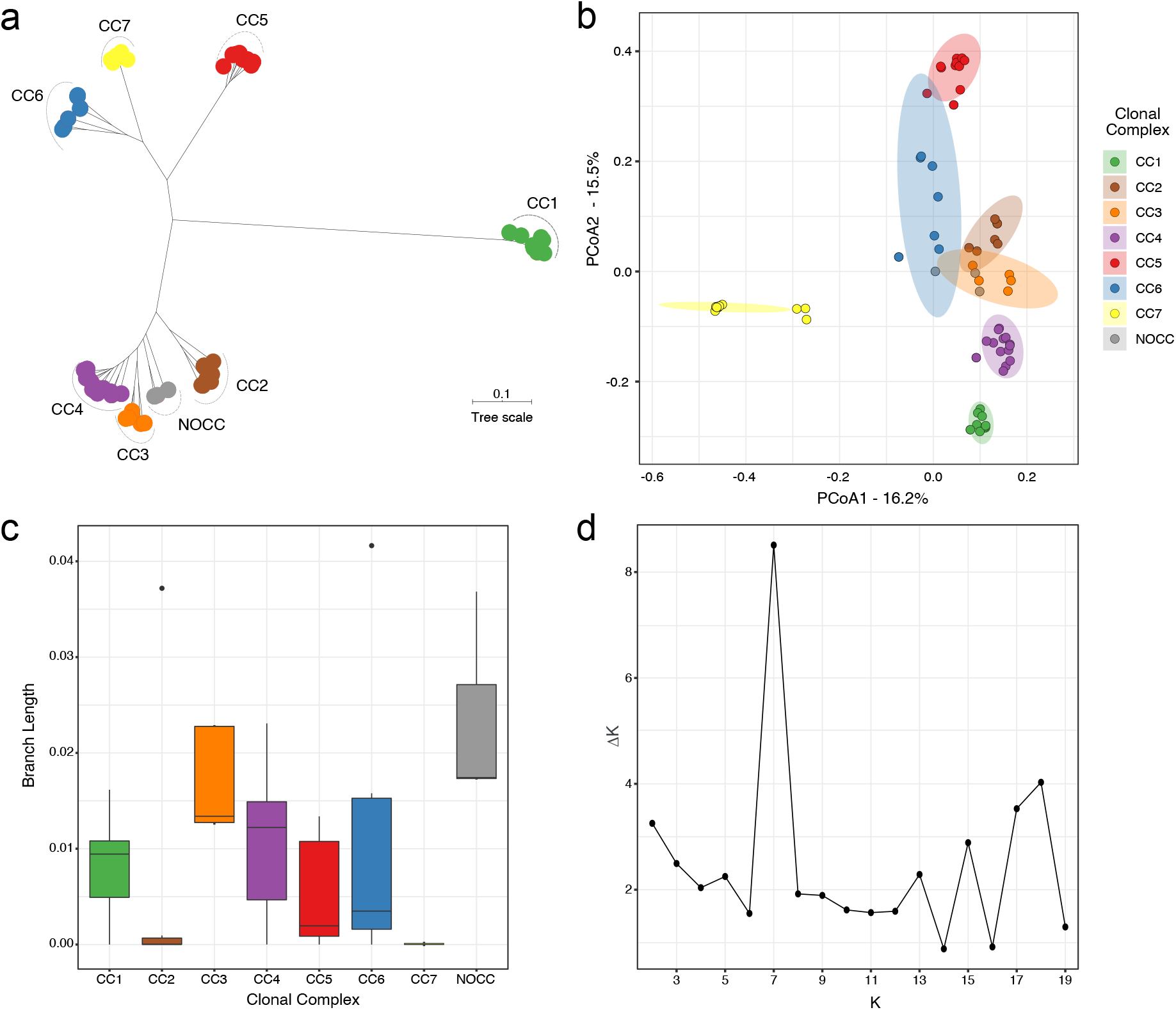
Population structure of *P. alloputida*. **a.** phylogenetic reconstruction using SNPs extracted from core genome to assign the cgMLST scheme. Colors represent distinct Clonal Complexes and NOCC stands for No Clonal Complex assigned with high confidence. **b.** Principal Component Analysis based on the presence/absence profile of accessory genes present in 5% to 95% of the isolates. **c.** Branch lengths for each Clonal Complex. **d.** ΔK distribution to estimate the best value for K, which supports the presence of 7 *P. alloputida* CCs.

We also performed a Discriminant Analysis of Principal Components (DAPC) [28] to recover the genes that contribute most to separate the population based on their presence/absence profiles (Figure S4). The top 50 discriminating genes separate CC5 and CC1 from the rest of the population, but fail to resolve the relationships between other CCs (Figure S5). Next, we used branch lengths from cgMLST tree as an indirect estimator of diversity within the CCs (Figure 3c). CC7 is the most clonal group, comprising 9 isolates, including KT2440 and three clinical isolates (GTC_16482, GTC_16473, and NBRC_111121).

The MLST scheme for *P. putida* species comprises eight housekeeping genes (*argS, gyrB, ileS, nuoC, ppsA, recA, rpoB*, and *rpoD*) [29]. We assigned isolates to STs and grouped those into CCs (Table S2). This analysis revealed some incongruences with previous reports [29]. For example, by using allele combinations with perfectmatch to predict STs, KT2440 was detected as ST69 and not as ST58 [29], indicating that a revision in the public database is warranted. The predicted number of CCs was also supported by the admixture model from STRUCTURE [30]. This model assumes that each isolate has ancestry from one or more *K* genetically different sources, which we referred to as CCs. The number of CCs corresponds to the estimated number of clusters represented by parameter *K*. Instead of using the highest raw marginal likelihood, we followed the protocol for estimating the best *K* value suggested by Evanno et al [31]. The *ad hoc* statistics *∆K* indicates that, according to our dataset, the population structure of *P. alloputida* is composed of at least 7 CCs (Figure 3d).

Once data from eight *loci* may be insufficient to accurately describe the population structure of *P. alloputida*, the populations identified by STRUCTURE were only considered if they matched cgMLST results. When correlating the cgMLST tree topology with ancestry proportion predicted for each isolate, we observed a clear delimitation of genetic blocks for each CC (Figure 4). However, there are a few inconsistencies. For example, isolate B4 has a greater ancestry proportion with CC2, although cgMLST indicates its greater proximity to CC3. In addition, KCJK7916, FF4, and B6-2 were not assigned to a CC because, although they have a higher proportion of ancestry with CC3, they are paraphyletic to CC3 and CC4. Since CCs are assumed to be monophyletic, these strains were designated as No Clonal Complex (NOCC). We expect that a greater number of *P. alloputida* genomes from isolates from various sources will improve the resolution of the *P. alloputida* population structure, including the CC assignment to isolates described here as NOCC.

**Figure 4.**
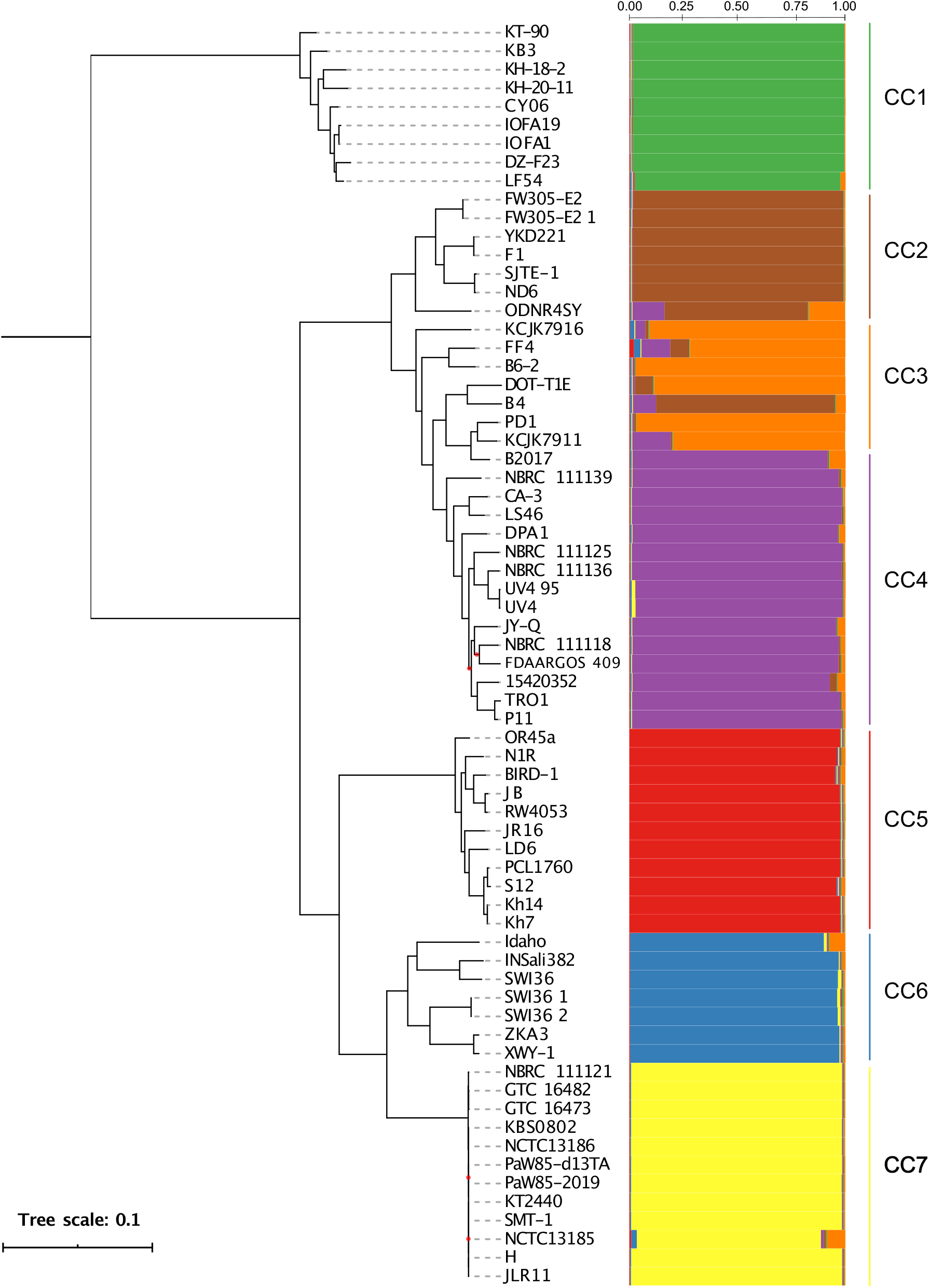
Phylogenetic tree and coancestry barplots correlating the phylogenetic tree using SNPs extracted from core genome and coancestry probabilities assigned with STRUCTURE. Red dots in branches represent bootstrap values lower than 70%. Each color represents one of the seven predicted Clonal Complexes (CC).

The cgMLST phylogenetic tree and the distance tree support CC1 as the basally branching group of *P. alloputida* (Figure 4). CC1 comprises isolates from deep-sea sediments from Indian Ocean, costal water from the Pacific Ocean, lotus field, and arthropods (Table 1). The main difference of CC1 is the lack of 229 gene families in the accessory genome, which are present in at least one isolate from all other CCs (Table S3). Among these absent gene families, there is a genomic island with nearly 46 Kbp encompassing 38 genes. Some of those genes are involved in sugar transport, as previously identified in KT2440 (CC7) [6], as well as genes encoding hypothetical proteins (coordinates 3,126,465-3,172,496 in KT2440).

**Table 1.**
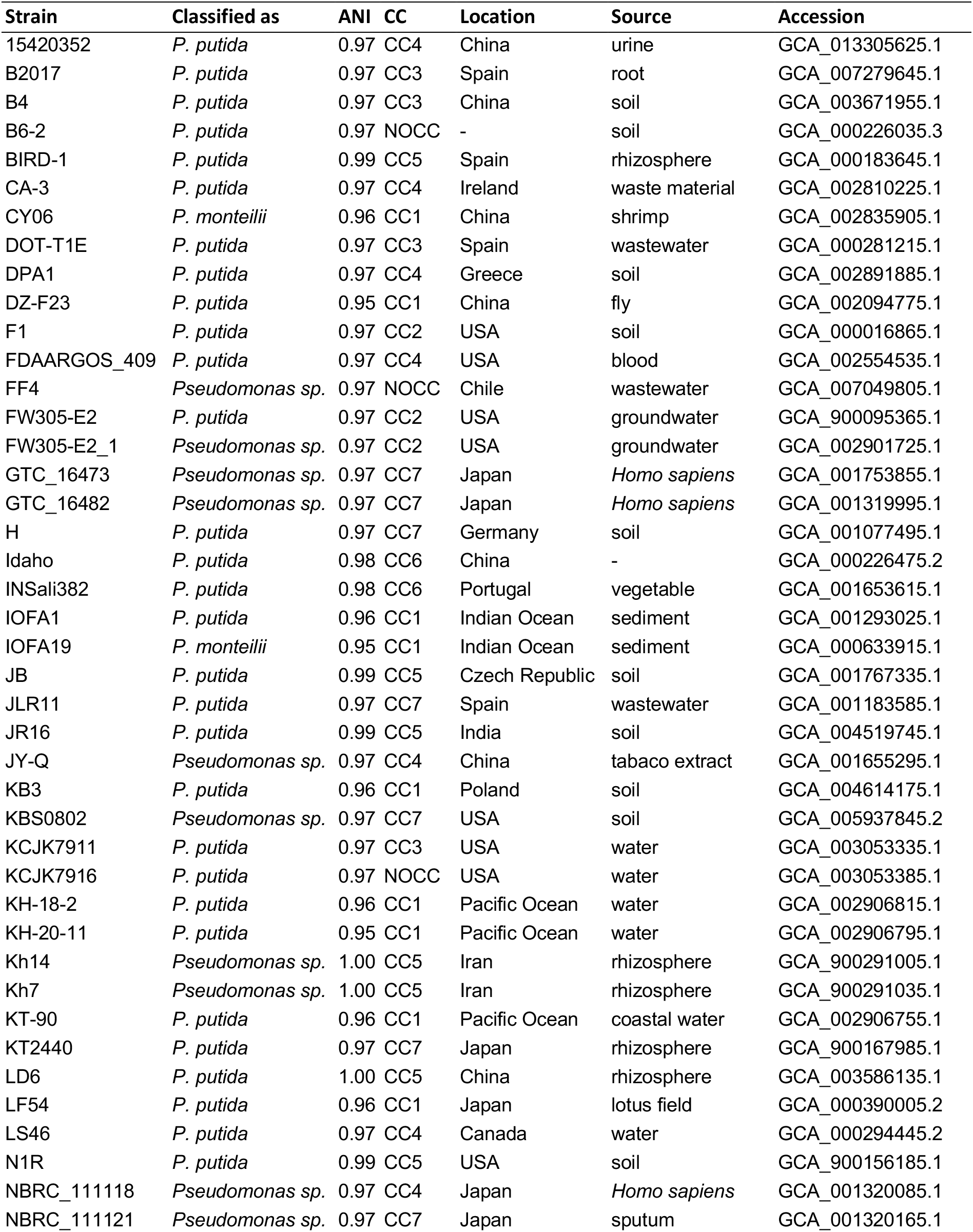
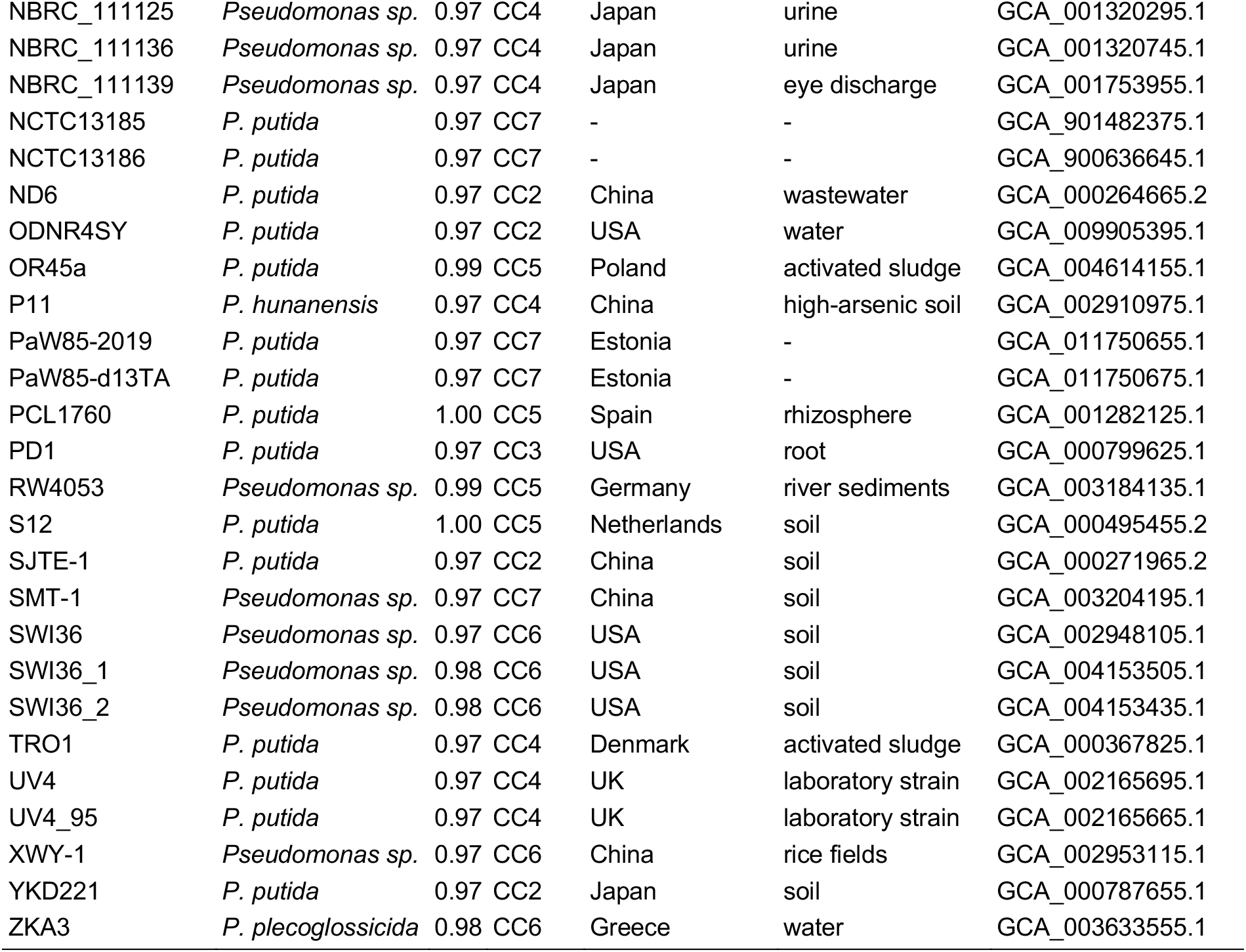
*Pseudomonas alloputida* isolates used in this study with average nucleotide identity (ANI) from Kh7^T^.

### 2.4. Resistance profiles

We evaluated the composition of antibiotic resistance genes using the CARD database [32]. All 15 different genes in the core resistome encode MDR efflux pumps (Table 2, Table S4), including MexAB-OprM, MexEF-OprN, and MexJK, from the resistance-nodulation-cell division (RND) efflux pumps family. These efflux pumps are associated with intrinsic and acquired multidrug resistance in *P. aeruginosa* [33, 34]. However, these RND efflux pumps may play an alternative role in *P. alloputida* by pumping out toxic substances such as toluene [35]. We also identified *cpxR*, which encodes a protein that promotes MexAB-OprM expression in the absence of the MexR repressor in *P. aeruginosa* [36], which is absent in *P. alloputida*. The presence of MexAB-OprM in the core genome, under CpxR regulation, supports its involvement with intrinsic physiology in addition to drug resistance, because this complex can be involved in both quorum-sensing and mediation of *P. aeruginosa*-host interaction [37, 38].

**Table 2.**
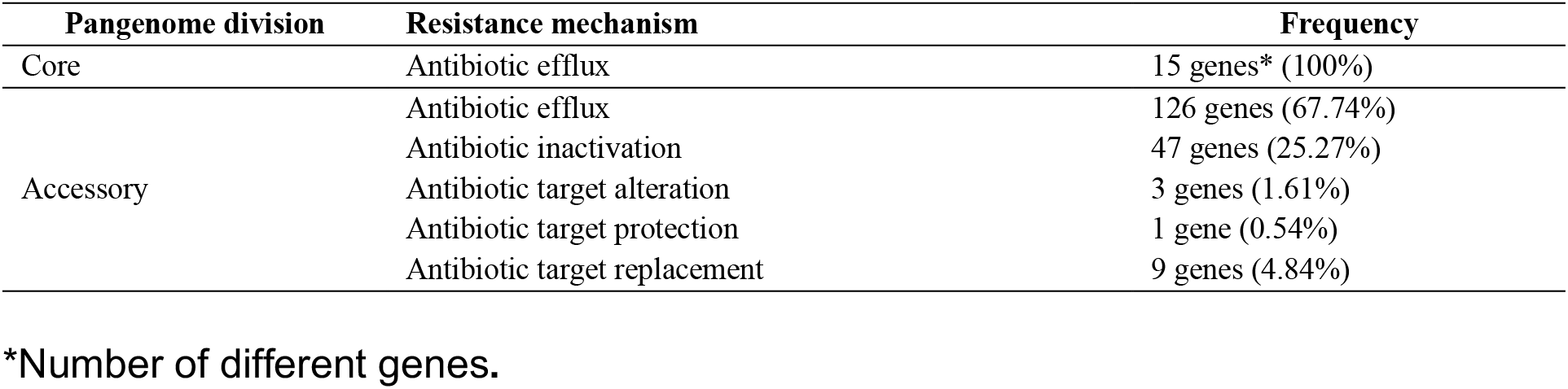
Frequency of resistance mechanisms categories in both core and accessory resistome.

Regarding the acquired resistome, we found 45 different genes that confer resistance by pumping out or inactivating antibiotics, as well as by interacting with antibiotic targets (Table S4). These genes are distributed at low-frequency (Figure 5a), indicating that most of them are strain-specific or acquired through horizontal gene transfer. In general, there is no clear correlation between acquired resistome and population structure (Figure S6), although CC7 has more acquired resistance genes than other CCs (Figure 5c, Figure S6).

**Figure 5.**
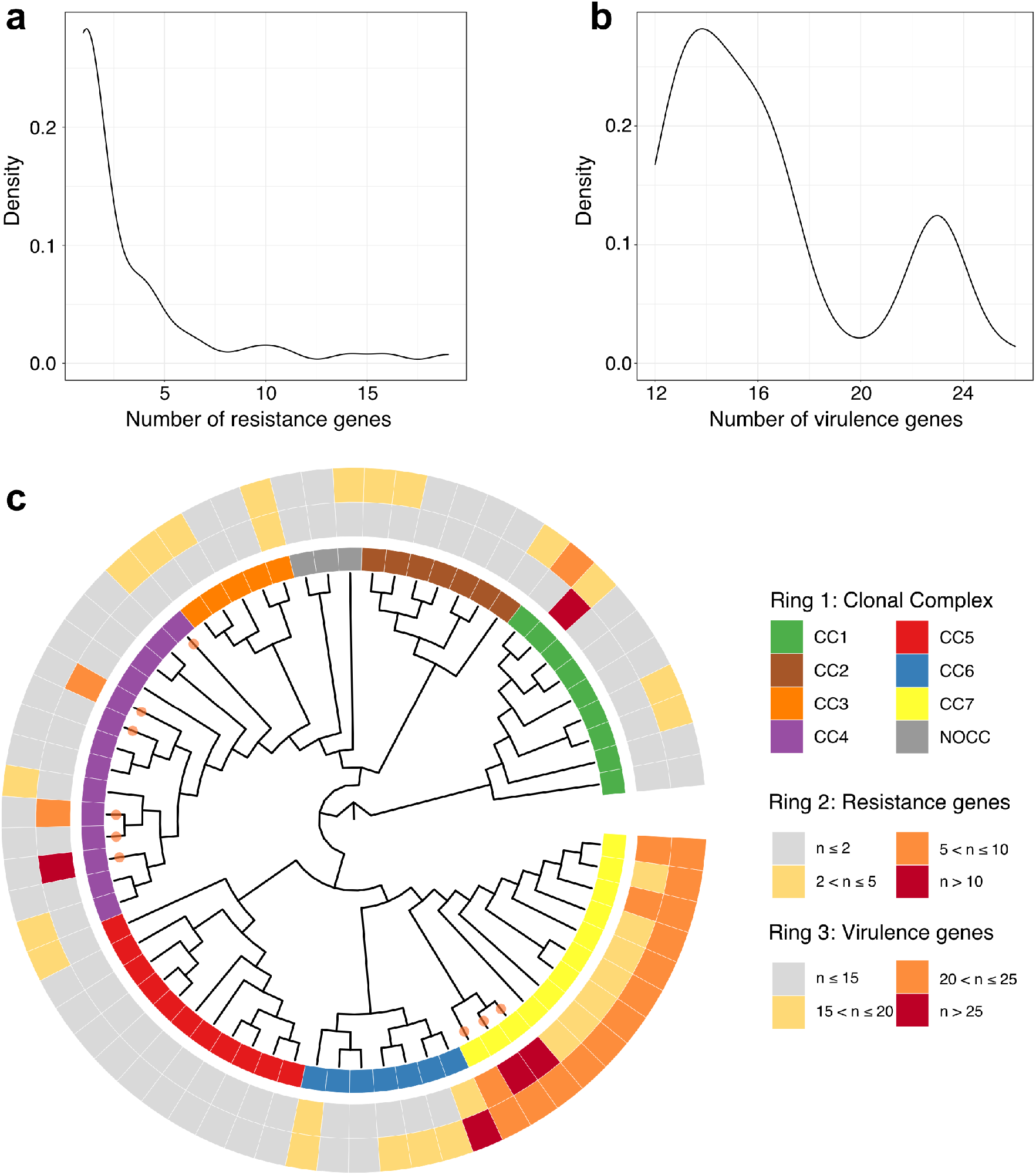
Population structure and distribution of virulence and resistance genes. **a, b.** distribution of acquired resistance and virulence genes. **c.** Maximum likelihood tree from SNPs present in the core genome inferred with 68 genomes used in this study. Branches from clinical isolates are marked with an orange circle. The inner ring represents the clonal complexes (NOCC: “No Clonal Complex”). The second and third rings indicate the number (n) of acquired resistance and virulence genes, respectively.

Our results highlight clinical strains harboring a range of resistance genes. In total, 9 out of 68 (13.2%) *P. alloputida* genomes analyzed here belong to clinical strains. Along with efflux pumps, the acquired resistome includes genes encoding antibiotic-inactivating enzymes that confer resistance to beta-lactams (e.g. *bla_cARB-3_*, *bla_IMP-1_*, *bla_oxA-2_*, *bla_PDC-7_*, *bla_TEM-1_*, and *bla_VIM-2_*); to aminoglycosides (e.g. *aac(6’)-IIa*, *aadA*, *aph(3”)-Ia*, and *aph(6’)-Id*; chloramphenicol (*cat*) and; to fosfomycin (*fosA*) (Table S2). These genes were distributed in few strains, mostly clinically relevant (Figure S6, Table S4); the top four strains with more acquired resistance genes were GTC_16473 (19 genes), GTC_16482 (16 genes), DZ-F23 (14 genes), and 15420352 (11 genes). Importantly, all of these strains (except DZ-F23) are clinical.

We evaluated the presence of acquired genes in plasmids predicted with the PLSDB database (version 2020_06_29) [39]. *P. alloputida* GTC_16473 contained the genes *aac(6’)-IIa*, *aadA23*, and *bla_cARB-3_* located in a scaffold with high identity with the pJR2 plasmid from *Pasteurella multocida* (NC_004772.1). We also identified *bla_OXA-2_*, *aadA22*, *aac(6’)-Ia*, *aac(6’)-IIc*, *aph(3”)-Ib*, *aph(6’)-Id*, *bla_IMP-1_*, and *sul1* genes in plasmid-like sequences in *P. alloputida* GTC_16473 that have not reached the coverage thresholds to be reliably classified as plasmids, supporting an underestimation of plasmids in *P. alloputida* isolates. *P. alloputida* XWY-1 (CC6) also contains a plasmid, pXWY-1 (NZ_CP026333.1), which harbors the resistance genes *sul1, aadA2,* and *qacH*. This strain was isolated from rice fields in China. Finding non-clinical strains harboring plasmids with such relevant resistance genes warrants further investigation.

### 2.5. Virulence profiles

We used the VFDB database [40] to assess the *P. alloputida* virulence profiles. Although strains such as KT2440 have been approved as a host-vector system safety level 1 [41], the genetic proximity with animal (*P. aeruginosa)* and plant (*P. syringae)* pathogens reinforces the need to evaluate the biosafety of *P. putida* species. The core virulome of *P. alloputida* contains genes associated with twitching motility, siderophore production (pyoverdine), and alginate biosynthesis (Table S5). Importantly, *P. alloputida* lacks key virulence genes usually found in *P. aeruginosa*, such as those encoding exotoxin A, alkaline protease, elastase, rhamnolipid biosynthesis pathway components, phospholipase C, plant cell wall-degrading enzymes, and type III secretion system. Although we have detected genes associated with the type III secretion system in both core (HopAJ2 and HopJ1) and acquired resistome (HopAN1), these genes were experimentally ruled out as virulence factors [42].

Most of the alginate regulatory and biosynthesis systems are present in the *P. alloputida* core virulome (Table S5). The overproduction of alginate plays an important role in inducing the mucoid morphotype in *P. aeruginosa* during chronic infection [43]. However, the transcriptional regulatory gene *algM* (*mucC*), associated with genotypic switching, is absent in *P. alloputida*. Since the lack of AlgM leads to the non-mucoid phenotype in *P. aeruginosa* [8], it can account for the non-mucoid phenotype in *P. alloputida* under no water stress conditions. Although the genotypic switching is unlikely to occur in *P. alloputida*, alginate production is important under water stress conditions by promoting biofilm development and protecting from desiccation [44].

The acquired virulome comprised genes for type II and VI secretions systems, adherence, and iron uptake (Table S5). While most isolates had a low frequency of resistance genes, the virulence factors displayed a bimodal distribution (Figure 5b), a pattern that has been previously observed for *Klebsiella aerogenes* [45]. Genes from the acinetobactin gene cluster and HSI-I type VI secretion system were differentially distributed across CCs (Figure 6a), although it remains unclear whether these patterns emerged mainly from gene gain or loss. In *Acinetobacter baumannii*, iron uptake is mainly performed by the siderophore acinetobactin [46], which is synthesized by the proteins encoded by the *bauABCDE* operon. We found this operon in *P. alloputida* strains within CC5, CC6, and CC7 (Figure 6a). Further, this operon was surrounded by genes coding for proteins from chemotaxis sensory transducer (PP_2599) and aminotransferases (PP_2588) families (Figure 6b). Siderophore-mediated iron acquisition has been investigated in *P. alloputida* KT2440 [47] (CC7), but the role played by *bauABCDE* in this species is yet to be elucidated.

**Figure 6.**
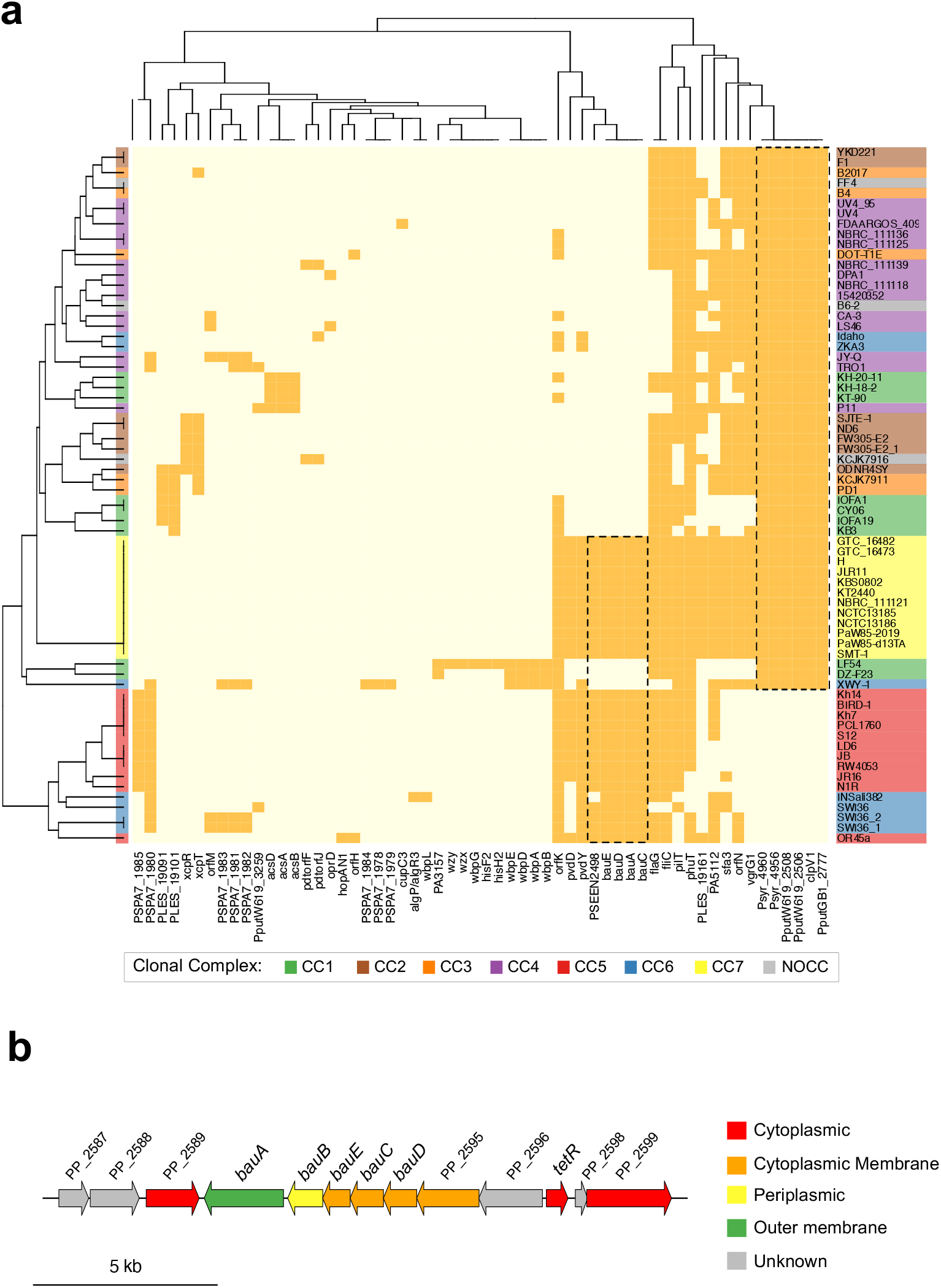
Acquired virulome composition. **a.** Matrix with presence (dark squares) and absence (light squares) profiles of virulence genes. Rows represent strains colored based on Clonal Complex that they belong. Columns are virulence genes identified. **b.** *P. alloputida* KT2440 *bauABCDE* genetic context annotated according Pseudomonas Genome Database (www.pseudomonas.com) and Song and Kim (2020).

Present in all strains from other CCs, the type VI secretion system (T6SS) HSI-I is absent in CC5 and CC6 (except XWY-1) isolates (Figure 6a). In KT2440 (CC7), HSI-I is a potent weapon against other bacteria (e.g. phytopathogens), increasing the competitiveness of *P. alloputida* [48]. The absence of T6SS virulence genes has been reported for BIRD-1 (CC5) [48], and our work generalizes this observation to all CC5 members. In KT2440, T6SS is crucial to kill phytopathogens such as *Xanthomonas campestris* [49]. These results indicate that BIRD-1 and other members from CC5 are likely less efficient than KT2440 as biocontrol agents. Moreover, all clinical *P. alloputida* strains harbor T6SS, indicating their potential ability to outcompete other bacteria during infections.

### 2.6. Plant growth promotion and bioremediation properties

The ability of *P. putida* species to promote plant growth and bioremediate toxic compounds have been explored [8, 10, 50, 51]. We searched for genes involved in plant growth promotion and bioremediation through a literature search of genes that have already been described for *Pseudomonas* species (Table S6). We found genes in the core genome, such as the pyrroloquinoline quinone-encoding operon *pqqBCDEFG,* associated with mineral phosphate solubilization in *Serratia marcescens* [52] and *Pseudomonas fluorescens* [53] (Table S7). Mineral solubilization has already been reported experimentally for BIRD-1 [50] and KT2440 [53], indicating that all *P. alloputida* isolates are genetically equipped to solubilize inorganic phosphate.

Another key feature that can enhance plant growth is the colonization of seeds by *P. putida* [54]. In KT2440, genes associated with surface adhesion (e.g. *lapA*, *lapBCD*), flagellum biosynthesis (e.g. *flhB*, *fliF*, *fliD*, *fliC*), and virulence regulation (e.g. *rpoN* and *gacS*) have been experimentally shown to be important for attachment to corn seeds [54]. All these genes, except *lapA* and *fliC*, belongs to the *P. alloputida* core genome (Table S7). Although often described as virulence genes, flagellum genes also play roles in the association of *P. putida* with plants. Other plant growth-promoting genes were also found in the *P. alloputida* accessory genome (Table S7), except for CC1, which lacks all putative plant growth-promoting genes.

Many plant growth-promoting bacteria can synthesize phytohormones such as indole-3-acetic acid. *P. alloputida* lacks the main genes involved in this process (e.g. *ipdC*, *iaaM,* and *iaaH*), indicating an incomplete or nonfunctional pathway. Although *P. putida* W619 (GCA_000019445.1) is considered one of the most efficient producers of IAA in comparison with other endophytic bacteria [19], this strain belongs to a different species. In addition, *P. alloputida* also lacks AcdS, an enzyme that counteract ethylene stress response, a result that has been experimentally confirmed in KT2440 [55].

Besides the ability to promote plant growth, *P. putida* can tolerate or degrade an array of compounds including heavy metals and hydrocarbons. We identified genes in the core genome that allow *P. alloputida* to resist various heavy metals such as copper (*cop* genes) and cobalt/zinc/cadmium (*czcABC*) (Table S8). The copper/silver resistance operon, *cusABC*, was present in the accessory genome. Although we found a wide range of genes associated with bioremediation, there is no clear correlation between the population structure and presence/absence profiles of such genes.

*P. putida* is known for its capacity to metabolize aromatic hydrocarbons such as toluene, benzene, and *p*-cymene [6, 8]. The toluene-degrading pathway includes the *todABCDE* operon and the *todST* regulator. The *p*-cymene compound can be degraded by means of the *cymAaAbBCDER* or the *cmtAaAbAcAdBCDEFGHI* operon [56]. We found a genomic island of approximately 48kb long harboring all these genes in F1, DOT-T1E, UV4, UV4/95, YKD221, and NBRC_111125 genomes (Figure S9). All these isolates, except NBRC_111125, were experimentally confirmed to degrade toluene. Further, *P. alloputida* F1 is well-known to grow on toluene [6]; YKD221 was isolated from contaminated industrial soil and degrades *cis*-dichloroethene [57]; DOT-T1E is an isolate known to grow on different carbon sources [58] and; UV4 and UV4/95 conduct important industrial biotransformation of arenes, alkenes, and phenols [59]. Interestingly, genes in this genomic island presented a very similar genetic context (Figure S7), with an upstream arm-type integrase associated with bacteriophages. We were unable to precisely define the *att* viral sites, indicating a deterioration of the original structure of the putative bacteriophage. Further, the lack of correspondence between population structure and the presence of an integrase upstream the genomic island indicates that this region was likely acquired via independent horizontal gene transfers in distinct *P. alloputida* CCs.

We also identified RND efflux pumps involved with solvent tolerance in both core (TtgABC) and accessory genomes (TtgDEF and TtgGHI). TtgABC, TtgDEF, and TtgGHI are required for DOT-T1E to efficiently tolerate toluene [35]. We observed that TtgABC is the same protein-complex predicted as MexAB-OprM, associated with antibiotic resistance in the core genome. This complex extrudes both antibiotics and solvents such as toluene in *P. alloputida* DOT-T1E [35], corroborating the additional and important function to extrude antibiotics and organic solvents in all *P. alloputida* isolates. TtgDEF is located in the same genomic island of *tod* genes. This complex can expel toluene, but not antibiotics [60], reinforcing the variety of molecules that can be extruded by RND efflux pumps and the need to explore the structural basis of this specificity, not only in *P. alloputida* isolates, but also in other bacteria.

## 3. Concluding remarks

Through a remarkable metabolic versatility, *P. putida* species can thrive in a wide variety of niches. In this work, we explored the genetic diversity of *P. alloputida* and characterized its population structure for the first time. Through a large-scale genomic analysis, we identified a major problem with *P. putida species* classification, including several reference strains that likely belong to new species, as also suggested elsewhere [13]. *P. alloputida* has an open pangenome dominated by low-frequency genes. The population structure of this species has at least 7 clonal complexes that were verified by cgMLST and STRUCTURE ancestry simulations. We found that most of the clinical isolates belong to CC4. Together, our results indicate that *P. alloputida* clinical isolates are mainly opportunistic and do not pose considerable health concerns.

We analyzed genes of clinical and biotechnological interest. *P. alloputida* has several RND-family efflux pumps that are important to tolerate antibiotics and other toxic compounds such as toluene. The low-frequency acquired resistance genes are predominant in plasmids from a few clinical strains. The detected virulence genes allow *P. alloputida* to synthesize pyoverdine, attach to seed surfaces, and kill other bacteria (including pathogens) through a type VI secretion system. KT2440 is a member of CC7 that has the T6SS along with an operon for acinetobactin biosynthesis. *P. alloputida* lacks key genes for the production of indole-3-acetic acid. We also observed that the genes for the degradation of some aromatic compounds, including toluene, were likely horizontally acquired. Our results provide an opportunity for the development of biotechnological applications as well as insights into the genomic diversity of the novel species *P. alloputida*.

## 4. Methods

### 4.1. Datasets and genomic features

We recovered 11,025 genomes from the *Pseudomonas* genus in June 2020. To assess the quality of the genomes, we used BUSCO v4.0.6 [14] with a minimum threshold of 90% completeness. The Kh7^T^ (GCA_900291035.1) was used as a reference with mash v.2.2.2 [61] to find genomes with distances up to 0.05. We used mashtree [15] to generate the distance tree. The ANI analysis was performed with pyani 0.2.10 [62]. Network analysis was conducted in R with the igraph package (https://igraph.org). We removed S12(GCA_000287915.1) and KT2440(GCA_000007565.2) because they were duplicated genomes. Type strains and accession numbers used to define clusters in the network analysis are available in the Table S1. Gene prediction in all isolates was conducted with prokka v1.12 [63] to avoid bias in the identification of protein families. Plasmids were analyzed with PLSD v2020_06_29 [39].

### 4.2. Pangenome characterization

We inferred the *P. alloputida* pangenome using Roary 3.13.0 [64], with a minimum threshold of 85% identity to cluster proteins. Core genes were defined as those present in more than 95% of the isolates. Jaccard distances were computed by using accessory genes with prevalence between 5% and 95%. Gene content variations between *P. alloputida* ecotypes were inferred with a discriminant analysis of principal components (DAPC) using the *ade4* and *adegenet* packages [28], retaining the 30 principal components and 3 discriminant functions. Pangenome openness and fluidity were conducted with micropan [65] with 500 and 1000 permutations, respectively.

### 4.3. Population structure analysis

We used *in-house* scripts to extract the genes present in all isolates, which were aligned with MAFFT v7.467 [66]. SNPs were retrieved with snp-sites v2.3.3 [67] and SNP alignment was used as input to RAxML v8 [68] to reconstruct the phylogenetic tree using the general time-reversible model and gamma correction. Since we used only variable sites as input, we used ASC_GTRGAMMA to correct ascertainment bias with the Paul Lewis correction. One thousand bootstrap replicates were generated to assess the significance of internal nodes. We inferred the cgMLST scheme using the core genome SNP phylogenetic tree. The phylogenetic tree was visualized with iTOL v4 [69].

We downloaded the *P. putida* MLST scheme (on June, 2020) containing 116 different STs (https://pubmlst.org/databases/). This scheme was designed for the whole *P. putida* group, not only *P. putida* sensu stricto [29]. We used BLASTN [70] to determine the best-matching MLST allele to access STs. The allelic profile associated with each ST in our dataset was used to conduct population assignment with STRUCTURE v2.3.4 [30] with admixture model. The length of Markov chain Monte Carlo (MCMC) was 50,000, discarding 20,000 iterations as burn-in. The simulations to calculate the parameter K ranged from 2 to 20, with 20 replicates for each K to estimate confidence intervals. Instead of using raw posterior probability to get the best K, we followed the protocol suggested by Evanno, Regnaut and Goudet [31]. Briefly, we calculated the first and second derivatives, resulting in a ΔK of 7. Therefore, we used K = 7 to analyze predicted ancestry probabilities.

### 4.4. Detection of genes associated with antimicrobial resistance, virulence, plant growth promotion, and bioremediation

We used the Comprehensive Antimicrobial Resistance Database (CARD) database v3.0.9 [32] to predict antibiotic resistance genes. The virulence factor database (VFDB) [40] was used to determine virulence genes. This database was downloaded on July 30 2020 and comprises 28,639 proteins associated with virulence in several pathogens. We used virulence genes previously described for the *Pseudomonas* genus. We clustered proteins based on 70% identity to build a non-redundant database using uclust v1.2.22q [71]. We built the database with plant growth promotion and bioremediation through literature searches (Table S6). All predicted proteins were globally aligned against these databases using usearch v11.0.667 [71] with 50% minimum coverage for query and subject and 60% minimum identity.

## Acknowledgements

This work was supported by Fundação Carlos Chagas Filho de Amparo à Pesquisa do Estado do Rio de Janeiro (FAPERJ; grants E-26/203.309/2016 and E-26/203.014/2018), Coordenação de Aperfeiçoamento de Pessoal de Nível Superior-Brasil (CAPES; Finance Code 001), and Conselho Nacional de Desenvolvimento Científico e Tecnológico. The funding agencies had no role in the design of the study and collection, analysis, and interpretation of data and in writing.

