## Supplementary Figures for "Phylogenetic and population structure analyses uncover pervasive misclassification and help assessing the biosafety of *Pseudomonas alloputida* for biotechnological applications"

*Corresponding authors

Short running title: Population structure of *Pseudomonas alloputida*

Av. Alberto Lamego 2000, P5 sala 217; Parque Califórnia
Campos dos Goytacazes, RJ, Brazil
CEP: 28013-602
HPA:; TMV:

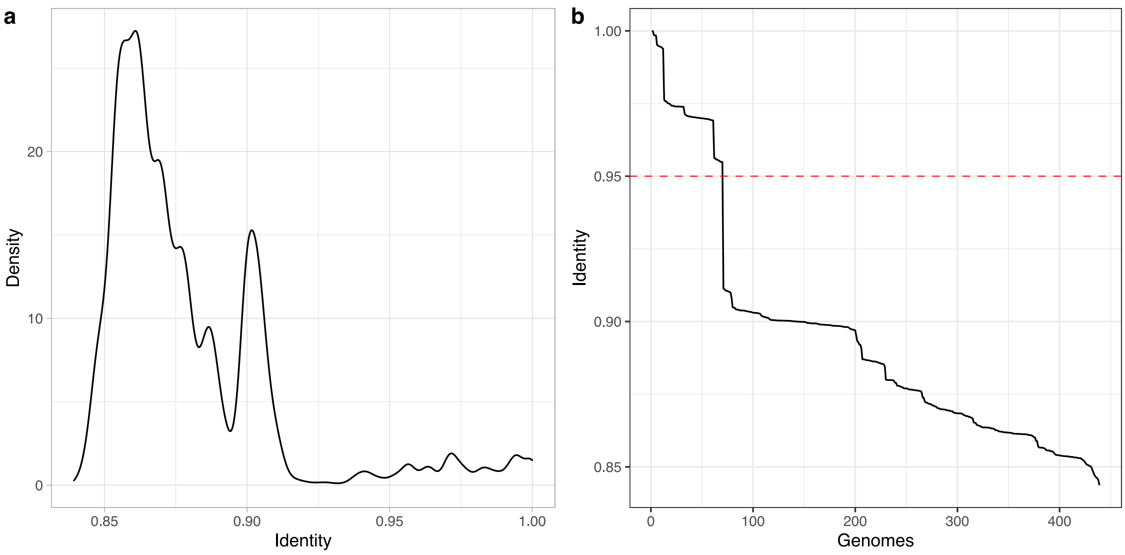


**Figure S1**. Average Nucleotide Identity distribution across *Pseudomonas putida* group. **a.** Identity density and **b.** ranked identity distribution in *P. putida* group from *P. putida* $Kh7^T$. Red dotted line represents the threshold used to define species based on average nucleotide identity.

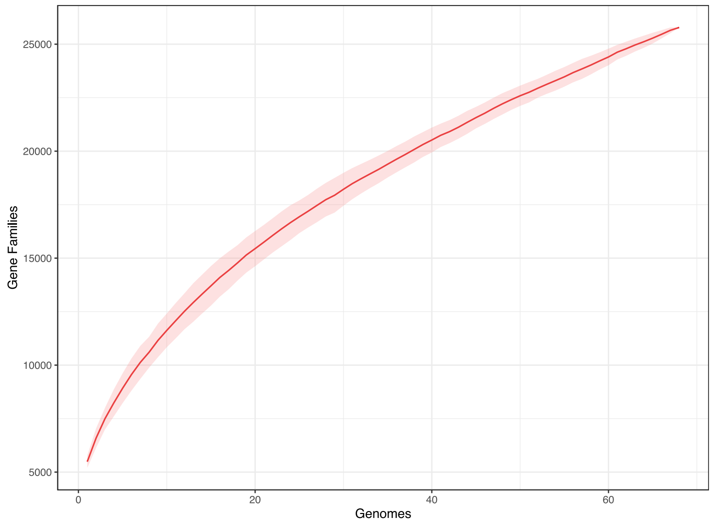


**Figure S2**. Cumulative curve of the *P. alloputida* pangenome. Gene families are in function of the number of isolates added sequentially. The slope ($\alpha$) of the curve is 0.417, indicating an open pangenome.

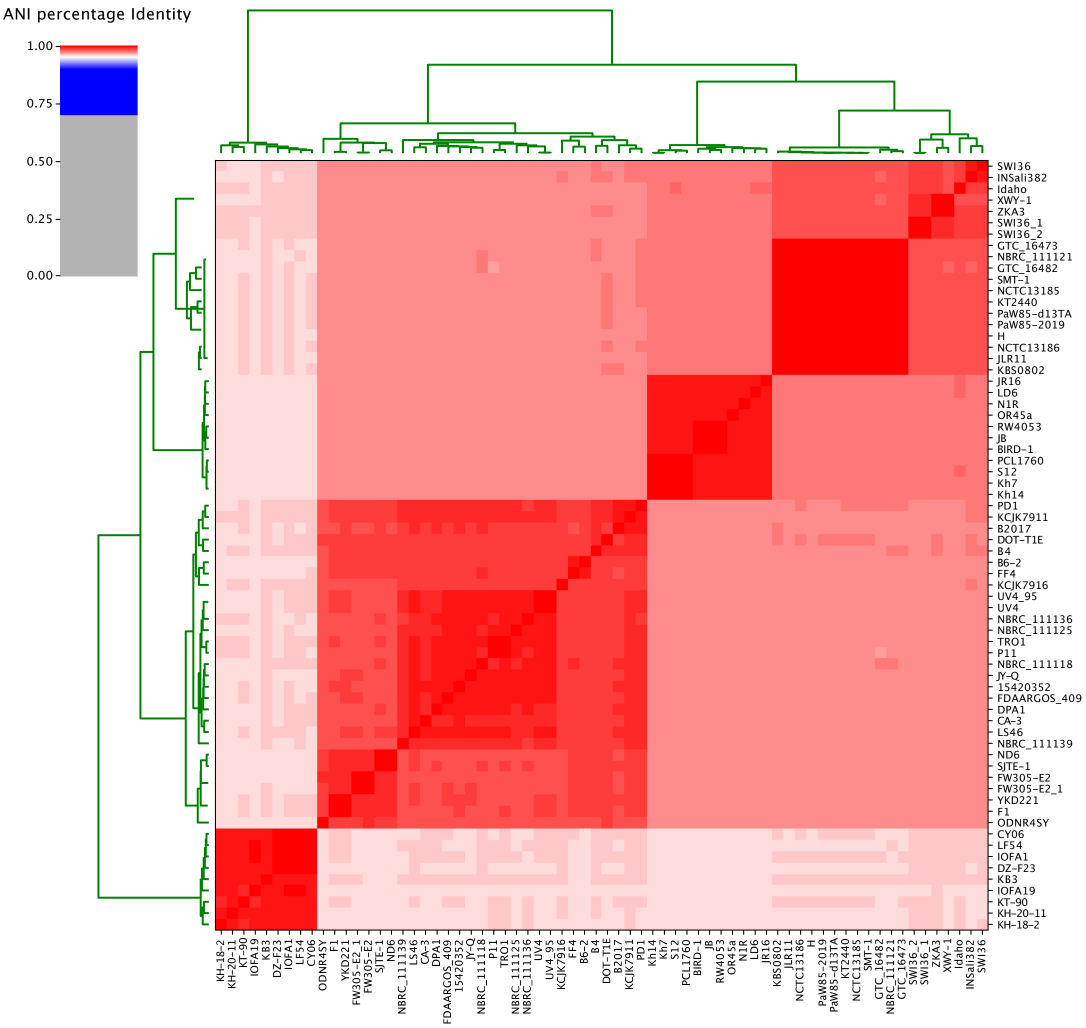


**Figure S3**. Average nucleotide identity analysis of the 68 isolates used in this study. Pairwise comparison indicating major groups in *P. alloputida*.

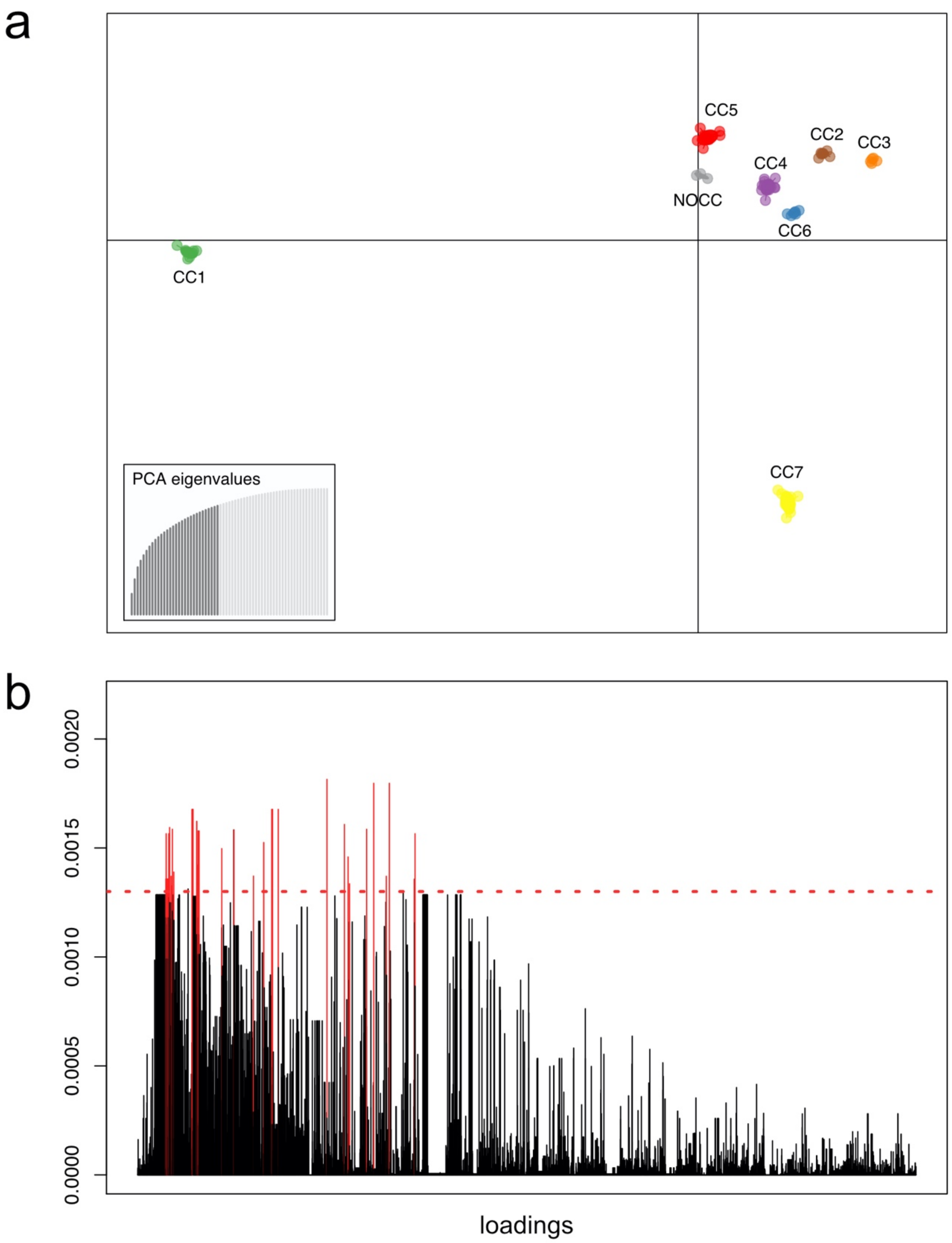


**Figure S4**. Discriminant Analysis of Principal Components of accessory genes present in 5% to 95% of the isolates. **a.** Clustering pattern of Clonal Complexes using the first two principal components of DAPC **b.** Loading plot. Red lines represent those gene families above the threshold of 0.0013 (red dotted line) and contributed more for the observed clustering patterns.

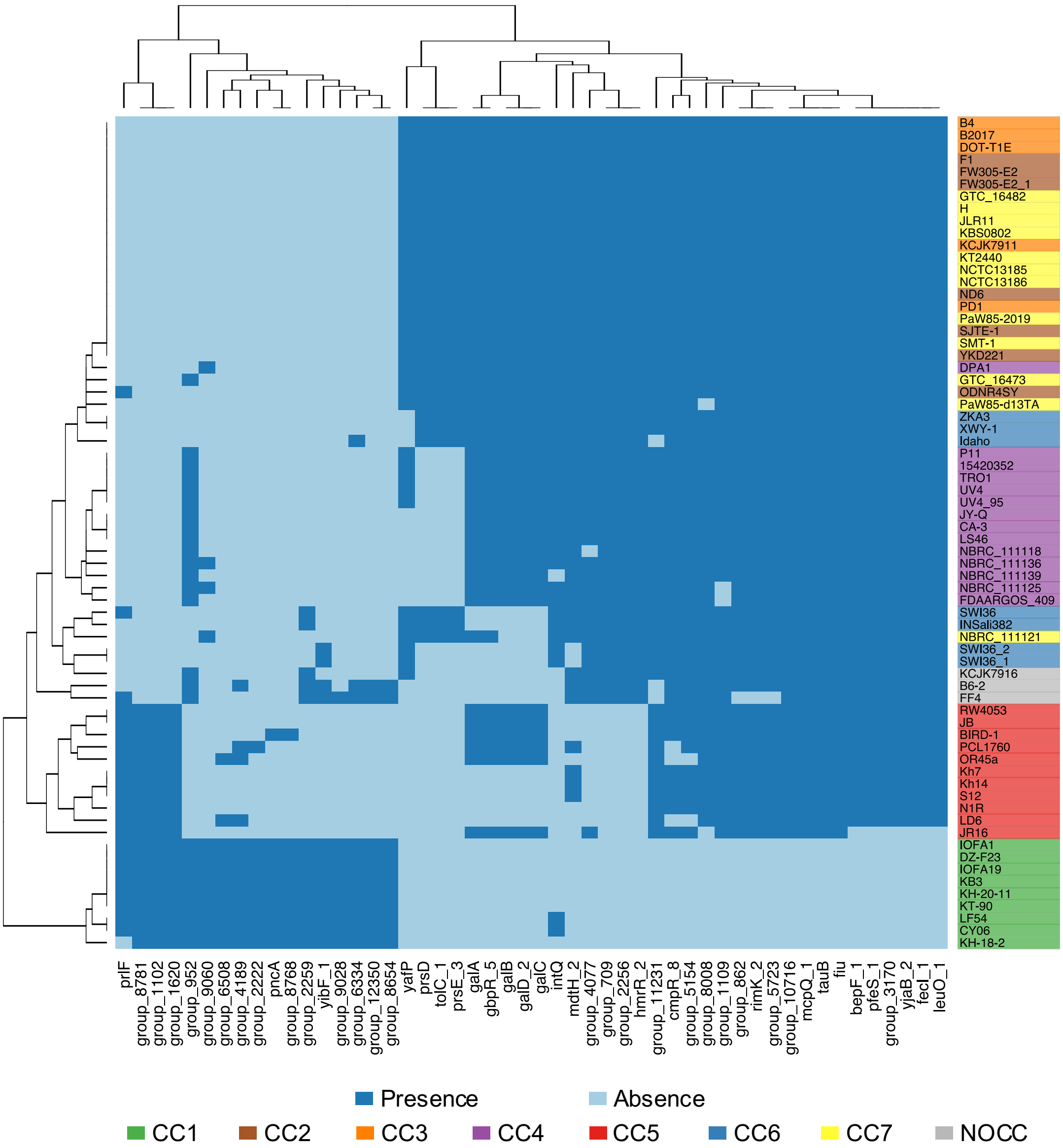


**Figure S5.** Heatmap from presence/absence profiles for top 50 genes detected to contribute for Clonal Complex clustering. Colors represent Clonal Complexes.

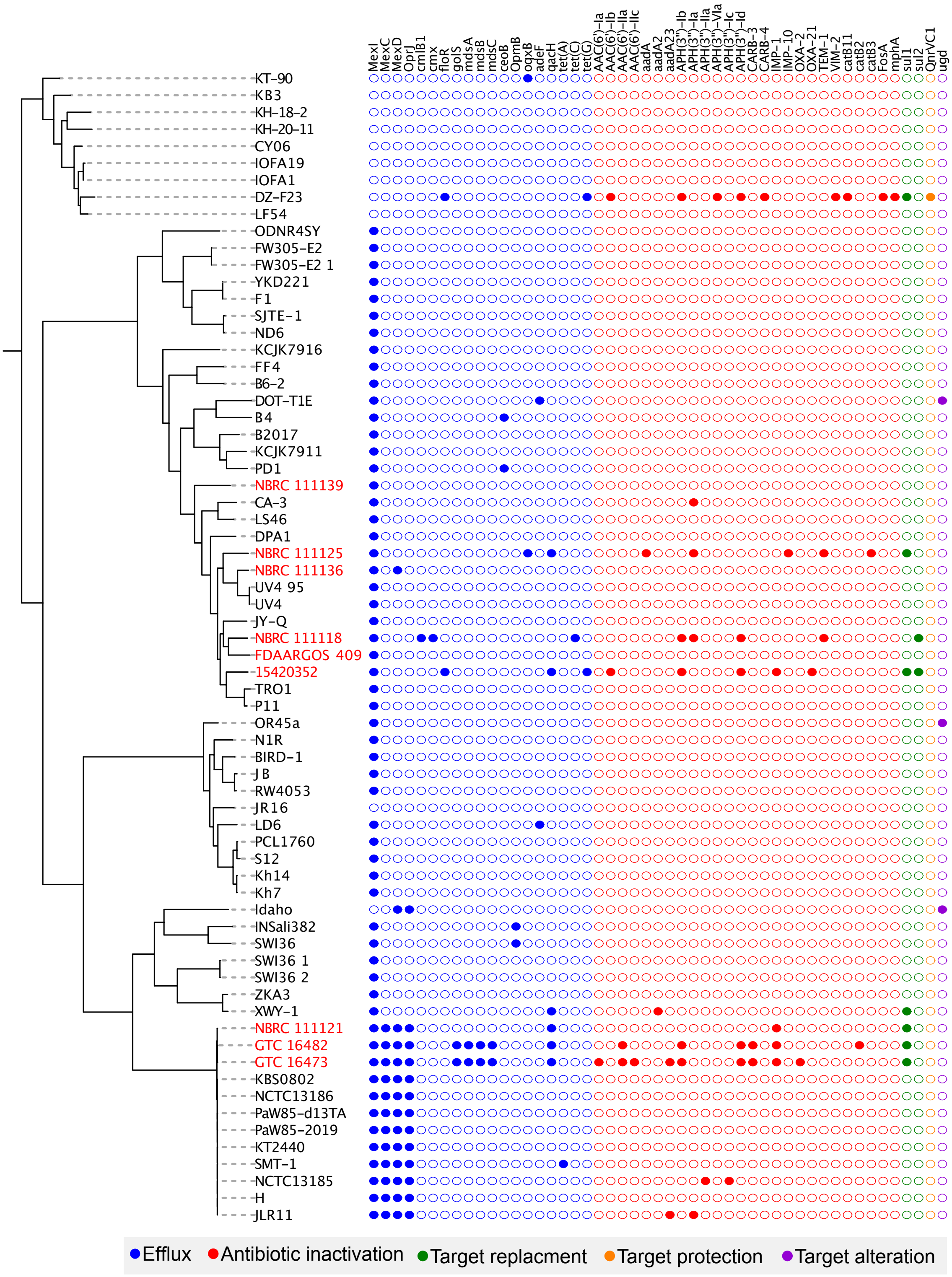


**Figure S6**. Accessory resistance genes identified in *P. alloputida*. Filled circles represent presence of a given gene and colors indicate the antibiotic resistance mechanism for each gene. Clinical strains are written in red.

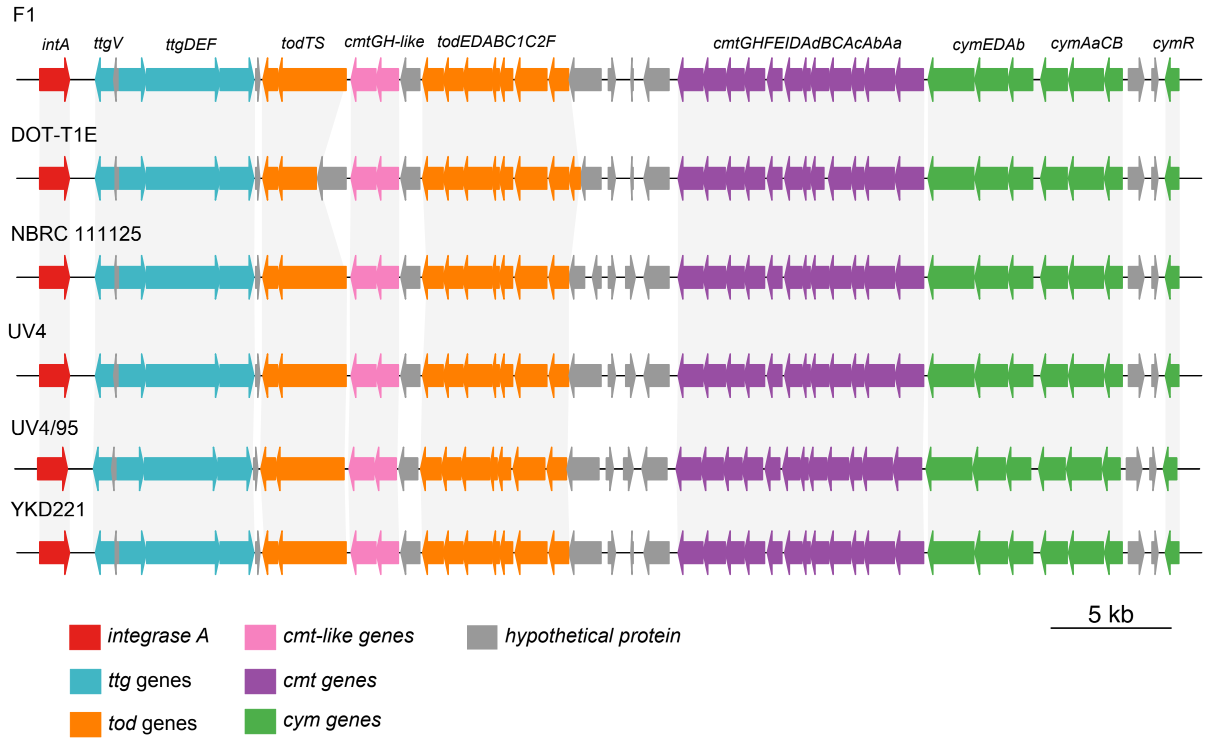


**Figure S7**. Genetic context of the genomic island containing *tod* genes in *P. alloputida* isolates. The gene coding for arm-type integrase A is upstream the genomic island, indicating a horizontal gene transfer for each represented isolate.
